# Targeted Ablation of Canonical Disulfide Bonds Mitigates TCR Mismatching in Engineered T Cells

**DOI:** 10.1101/2025.05.28.656745

**Authors:** Dan Su, Guangna Liu

## Abstract

T cell receptor-engineered T (TCR-T) cell therapy holds distinct advantages over chimeric antigen receptor (CAR)-T cell therapy in treating solid tumors by exploiting intracellular targets through MHC-mediated antigen presentation. Despite this potential, only one TCR-T therapy has received FDA approval to date. A key challenge lies in chain mispairing between exogenous therapeutic TCR chains and endogenous TCR chains, which compromises membrane localization efficiency and elevates off-target toxicity risks. While multiple engineering strategies have been developed to enhance TCR surface expression, our systematic evaluation using a novel mispairing assessment platform revealed that conventional approaches may paradoxically exacerbate chain mismatching. Through targeted cysteine mutations that disrupt the native interchain disulfide bonds in TCR constant domains, we achieved a substantial reduction in exogenous-endogenous TCR mispairing. Importantly, these structural modifications preserved both high surface expression level and potent tumor-specific cytotoxicity. This innovative engineering approach establishes a critical foundation for clinical-grade manufacturing of TCR-T therapies with enhanced safety profiles.

## Introduction

T cell receptor-engineered T cell (TCR-T) therapy involves genetic modification of human T cells with exogenous TCR genes to redirect their antigen specificity against tumor cells(1). Unlike chimeric antigen receptor (CAR)-T cells, TCR-T cells recognize antigenic peptides presented by MHC molecules, enabling targeting of both intracellular and extracellular tumor antigens(2). This capability positions TCR-T therapy as a promising approach for treating solid tumors(3), with clinical trials targeting diverse antigens(4) including NY-ESO-1 in synovial sarcoma(5, 6), melanoma(7, 8), and myeloma(7, 9).

Despite its potential, TCR-T development faces unique challenges. While antigen escape(10), on-target/off-tumor toxicity(11), and immunosuppressive microenvironments(12) are shared with other cell therapies, TCR-T faces an additional structural constraint: its heterodimeric αβ TCR chains must pair precisely to avoid mismatches with endogenous TCR chains(13, 14). The structural similarity between exogenous and endogenous TCR chains allows four possible heterodimer combinations during T-cell engineering(15). Mismatched TCRs—unlike thymus-selected endogenous TCRs(16)—may exhibit autoreactivity or alloreactivity(17), potentially causing off-target toxicity or graft-versus-host disease (GVHD)(18). Furthermore, competition between mismatched TCRs and therapeutic TCRs for CD3 complex binding reduces the surface expression of functional TCRs(19), impairing tumor-killing efficacy. Although GVHD has only been observed preclinically, eliminating TCR mismatching remains critical for clinical safety and potency(20, 21).

Current strategies to enhance TCR pairing efficiency include: (a) replacing human TCR constant regions with murine counterparts(22-24); (b) introducing an interchain disulfide bond (α48/β57)(23, 25); (c) modifying α-chain transmembrane domains with hydrophobic residues(26), and (d) incorporating murine-derived stabilizing mutations(27). While murine constant region substitution remains predominant in clinical studies, prolonged expression of xenogeneic sequences raises immunogenicity concerns. Moreover, the combinatorial effects of these strategies on eliminating exogenous-endogenous TCR mismatches remain unverified. Addressing these limitations requires innovative engineering approaches that minimize non-human sequences while achieving high-fidelity TCR pairing.

## Methods

### Plasmid Construction

The mutated TCR gene was cloned using the Phanta Max Super-Fidelity DNA Polymerase kit (Vazymes), the fragment was identified by DNA gel electrophoresis, and the gel was recovered using the Agarose Gel DNA Recovery Kit (TIANGEN). The lentiviral vectors containing the red fluorescent protein (RFP) genes were double cleaved by selecting the appropriate cleavage site, and the vector and gene fragment were homologously recombined using the pEASY* -Basic Seamless Cloning and Assembly kit (TransGen). After successful plasmid construction, plasmid extraction was performed using High Purity Plasmid Extraction kit (TIANGEN).

### Lentiviral Packaging

A four-plasmid system was used for lentiviral packaging. The target plasmid, pMD2.G, pRSV-Rev, and pMDLg were mixed at a 4:1:1:2 mass ratio in DMEM medium. PEI was used as a transfection reagent. The plasmids and PEI were mixed to form a transfection complex, which was then added to the L293T cells with a growth density of about 80%. The lentiviral supernatant was collected at 48 and 72 hours after transfection. The lentiviral supernatant was concentrated by PEG8000 then resuspended in serum-free 1640 medium and stored at -80°C.

### TCR-T Cell Construction

For experiments performed in Jurkat cells, wild-type and TCRαβ-KO Jurkat cells were seeded in 12-well plates at 5×10^5^ cells/well. Following the addition of 10 μL viral concentrate per well, cells were cultured for 13–14 hours before medium replacement with fresh medium. Transduced cells were then expanded for subsequent surface expression analysis and functional activation assays.

For experiments performed in human primary T cells, human peripheral blood mononuclear cells (PBMCs) were cultured in 1640 medium (Gibco, #C11875500BT) containing 10% fetal bovine serum (FBS) and IL-2 at 200IU/mL. Before infection, PBMCs were activated in a 48-well plate precoated with OKT3 (5 µg/mL), antibodies and RetroNectin (5 µg/mL, Takara). After stimulation for 24 hours, PBMCs were infected with viral particles at a multiplicity of infection (MOI) of 10-20. Then T cells were expanded for a total of 7–9 days before further assessment.

### Detection of TCR Surface Expression

To detect TCR surface expression, tetramer-APC diluted in 1×PBS/1% FBS/0.5 mM EDTA was added to cells, after incubated for 30 min on ice, cells were washed twice by 1×PBS then suspended in fixative solution(Solarbio, #P1110). Flow cytometry data were acquired by BD FACSCelesta. The surface expression efficiency of TCRs was calculated as follows: the RFP-positive cell population was gated, which represents the infection efficiency, then Tetramer-APC-positive cell population was gated from RFP^+^ population, which represents the surface expression efficiency.

### Jurkat Cell Activation Assay

T2 target cells were seeded in plates, and NY-ESO-1 peptide was added to a final concentration of 0.1 μg/mL. The cell-peptide mixture was then incubated for 2 hours at 37°C. TCR-transduced or control Jurkat cells were co-cultured with peptide-pulsed T2 cells or control T2 cells for 12 hours at an effector: target ratio of 1:4, then activation marker CD69 was detected by flow cytometry.

### Cytotoxicity assay

Engineered human primary T cells and Huh7-0201-NYESO1-luc/GFP target cells were co-cultured for 24 hours at an effector: target ratio of 1:10, and the luciferase activity of alive target cells was detected by Luciferase Reporter Gene Assay Kit (YEASEN). The bioluminescence signal intensity was detected by Microplate Reader (Agilent, Synergy H1). Then the percentage of target cell death was calculated.

### Data analysis

Data analysis in this study employed: SnapGene 6.1.1 for primer design and plasmid sequence alignment, FlowJo 10.8.1 for flow cytometry analysis and visualization, GraphPad Prism 10.0.2 for cytotoxicity assessment and cytokine quantification analysis. Statistical comparisons between different groups were determined by two-way ANOVA, and significant differences were indicated by *, * means p<0.05, ** means p<0.01, *** means p<0.001, **** means p<0.0001, and ns means no significant difference.

## Results

### Combinatorially Optimized TCR Exhibits High Level of Mismatch

To systematically evaluate the impact of reported TCR optimization strategies on chain mispairing, we aimed to design a combinatorial TCR construct that integrates multiple engineering approaches, which can possess high surface expression and cytotoxicity while minimizing non-human sequences. Previously reported strategies to enhance TCR pairing include: (a) replacing human TCR constant regions with murine counterparts; (b) introducing an extra interchain disulfide bond (α48/β57, termed 4857 in this study); (c) modifying α-chain transmembrane domains with hydrophobic residues substitution (referred as TMa in this study); and (d) incorporating murine-derived site mutations (referred as mm in this study). We used the well-known HLA-A*02:01-restricted 1G4 TCR as study model. Given that complete murine constant region replacement heightens immunogenicity risks, we implemented strategies b, c, and d on 1G4 TCR while retaining human constant regions, creating an optimized TCR (sTCR) (Figure 1A, 1B). Compared to TCR with fully human constant regions (hTCR), sTCR demonstrated dramatically enhanced surface expression (91.2% vs 36.8%, Figure 1C). To assess functional efficacy, we engineered Huh7 cells co-expressing NY-ESO-1 and HLA-A*02:01, along with stable luciferase/GFP expression (Figure 1D). When co-cultured with Huh7-A02-NY-ESO-1 cells, sTCR-transduced T cells exhibited superior cytotoxicity compared to hTCR-T cells (Figure 1E).

**Figure 1.**
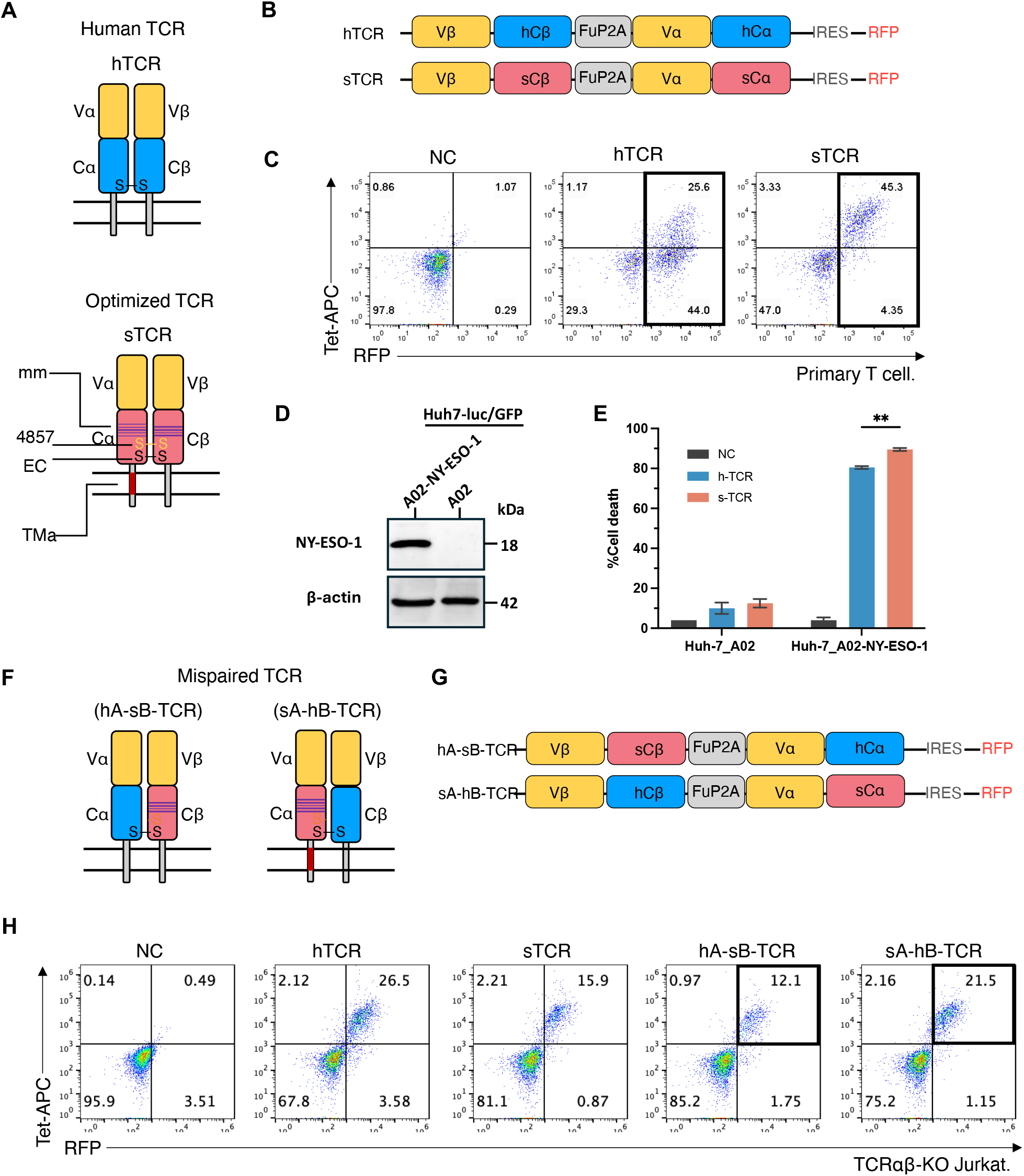
sTCR showed superior cytotoxicity but a high level of mismatch. (A) Schematic of hTCR and sTCR structures. (B) Schematic of hTCR and sTCR plasmids sequence. (C) Flow cytometry detection of hTCR and sTCR of the surface expression level in human primary T cells. (D) Western blotting assay to detect NY-ESO-1 protein in target cells. (E) Co-culture assay to detect the cytotoxicity of hTCR and sTCR, T cells and tumor cells were co-cultured for 24 hours at a effector:target ratio of 1:10,. Data represent the mean with SD of three technical replicates, statistical significance was determined by two-way ANOVA. ** means p < 0.01. (F) Schematic structures of the two mismatch constructs: hA-sB-TCR and sA-hB-TCR. (G) Schematic diagram of the plasmid sequences of the two mismatch constructs hA-sB-TCR and sA-hB-TCR. (H) Surface expression detection of the mismatch hA-sB-TCR and sA-hB-TCR in TCRαβ-KO Jurkat cells.

To specifically evaluate exogenous-endogenous TCR mispairing, we developed artificial mismatch detection constructs (Figure 1F, 1G). Upon transduction into TCRαβ-KO Jurkat cells, both sTCR chains showed substantial mispairing propensity with endogenous human TCR chains (Figure 1H). These results reveal a critical paradox: while combinatorial optimization enhances surface expression and cytotoxicity, it fails to address chain mispairing issue and the associated clinical safety risks in therapeutic applications.

### Disruption of Canonical Disulfide Bond Maintains Full Functionality in Engineered TCRs

The conserved αCys95–βCys131 disulfide bond represents the only known mechanism mediating chain pairing in endogenous human TCRs. Given that transduced therapeutic TCRs retain these residues, whether this intrinsic disulfide bond drives exogenous-endogenous TCR mispairing remains unexplored despite its fundamental role in natural TCR assembly. Thus, we engineered double cysteine mutations (αC95A/βC131A) in sTCR constant domains to disrupt this essential interchain disulfide bond (Figure 2A, B). Remarkably, the cysteine-mutated variant (mutEC-sTCR) maintained structural integrity with >95% antigen-specific binding (Tetramer-APC^+^/RFP^+^) in Jurkat cells (Figure 2C), demonstrating for the first time that functional TCR pairing and membrane localization can occur independently of this canonical disulfide bridge. In primary T cells, mutEC-sTCR showed comparable surface expression to sTCR (89% vs 80%, within 10% variation) (Figure 2D). Upon stimulation with peptide-pulsed T2 cells, mutEC-sTCR mediated enhanced activation compared to both hTCR and sTCR (Figure 2E). And the cytotoxicity of mutEC-sTCR-expressing T cells also outperformed hTCR-T cells while matching sTCR-T cells (Figure 2F). This confirms that engineered TCRs can bypass the conserved disulfide constraint while preserving superior functionality.

**Figure 2.**
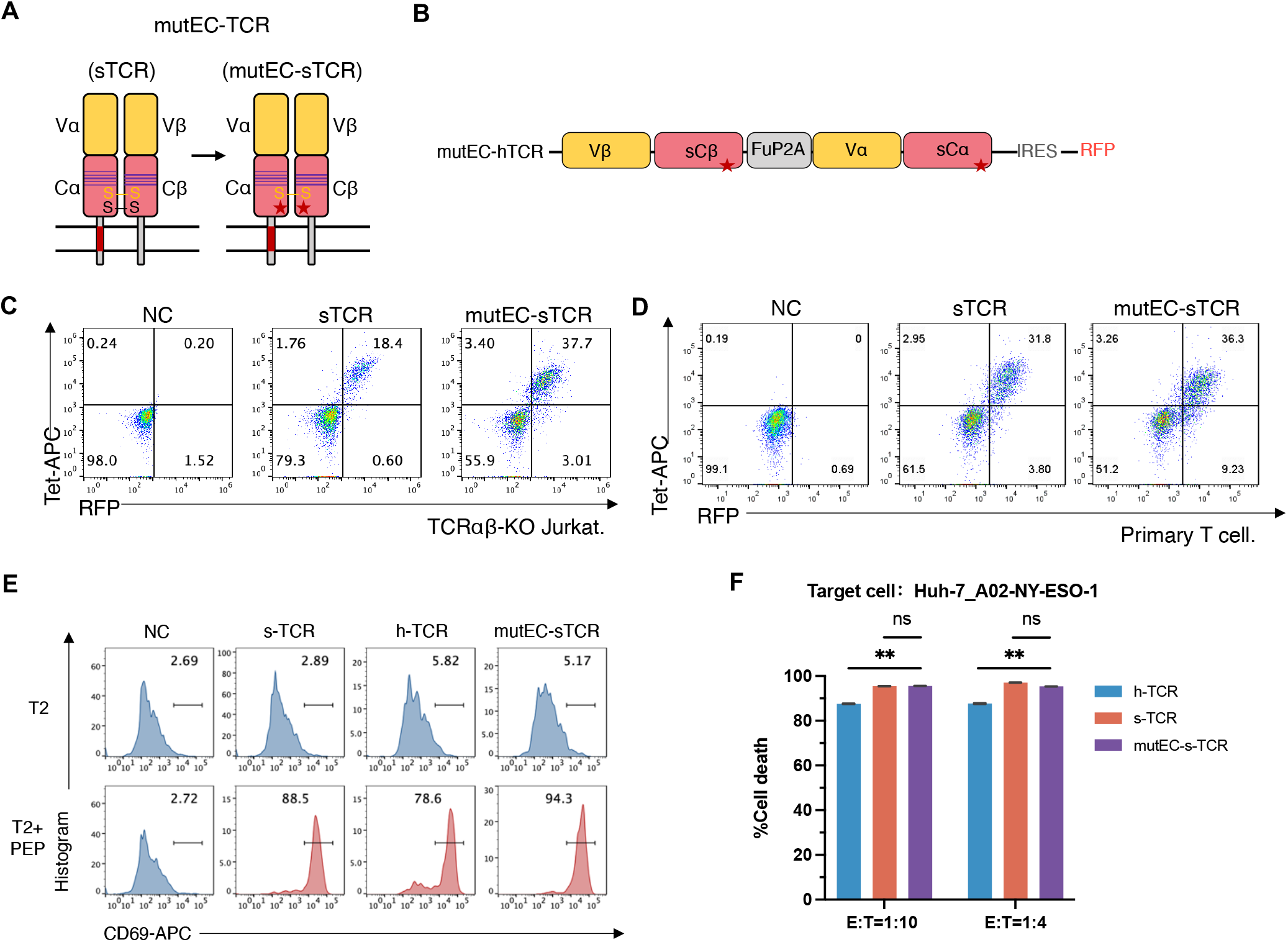
Disruption of canonical disulfide bond of sTCR does not affect its function. (A) Schematic of sTCR and mutEC-sTCR structures. (B) Schematic of sTCR and mutEC-sTCR plasmids sequence. (C) Flow cytometry detection of sTCR and mutEC-sTCR of the surface expression level in TCRαβ-KO Jurkat cells. (D) Flow cytometry detection of sTCR and mutEC-sTCR of the surface expression level in human primary T cells. (E) mutEC-sTCR-engineered Jurkat cells were co-cultured with T2 target cells pulsed with NY-ESO-1 peptide for 12 hours., and CD69 expression was detected by Flow cytometry. (F) mutEC-sTCR engineered human primary T cells were co-cultured with Huh7-A02-NY-ESO-1-luc/GFP cells and subjected to cytotoxicity assay after co-culture for 24h, effector: target (E:T) ratio was 1:10. Data represent the mean with SD of three technical replicates, statistical significance was determined by ordinary one-way ANOVA. ns means not significant, ** means p < 0.01.

### TCR Mismatch is Attenuated by Canonical Disulfide Bond Ablation

To further determine whether exogenous-endogenous TCR mispairing can be eliminated by ablation of the canonical disulfide bond, we reconstituted artificial mismatch detection constructs pairing hTCRs with mutEC-sTCR chains (Figure 3A, B). Assessment in TCRαβ-KO Jurkat cells revealed significantly reduced surface expression across these mutated configurations compared to original sTCR mismatch constructs (Figure 3C, Figure 1H). Notably, mispairing between the disulfide bond-deficient β chain (mutsB) and endogenous α chain was nearly abolished (Figure 3C).

**Figure 3.**
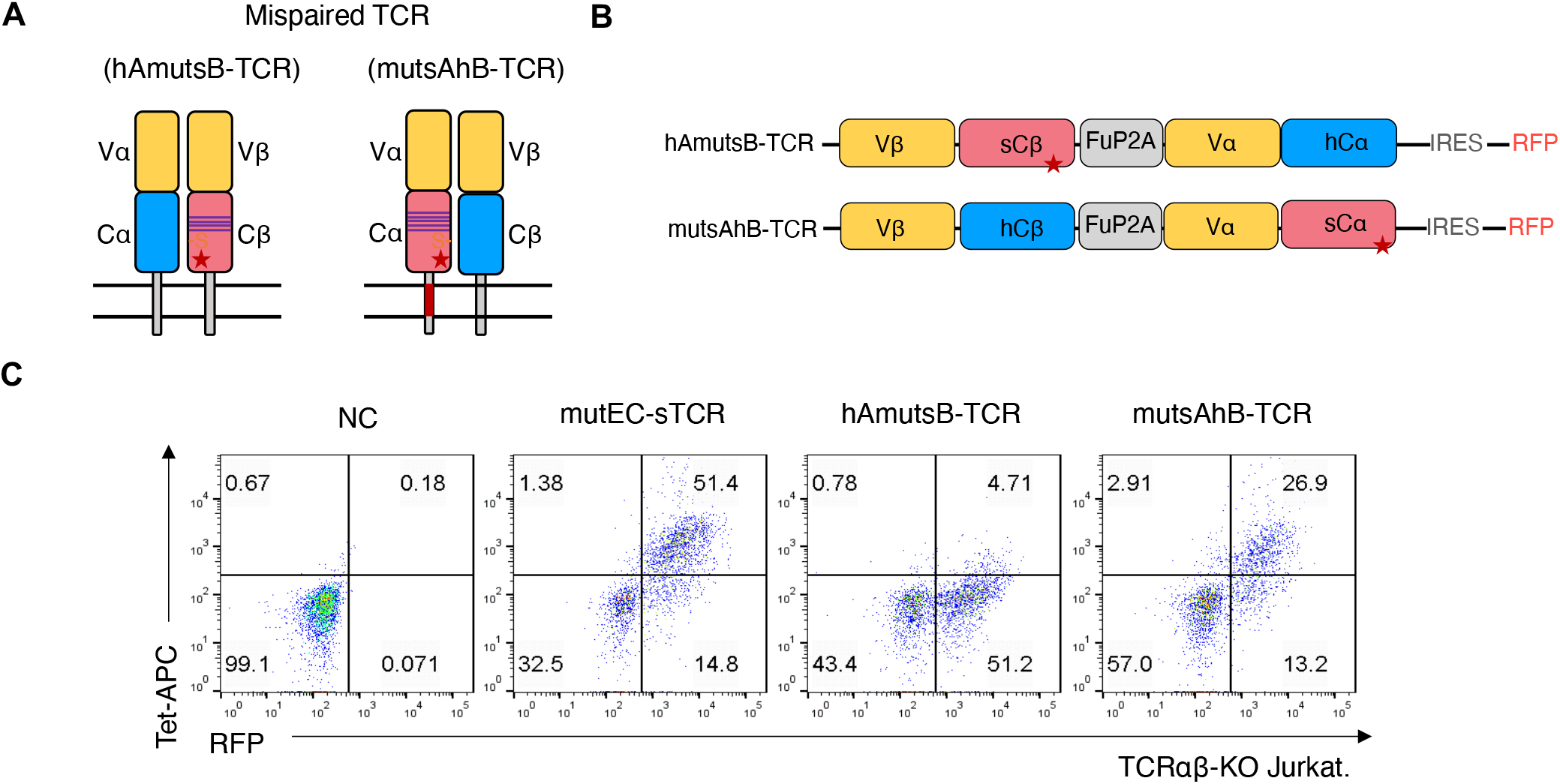
Canonical disulfide bond ablation reduces TCR mismatch. (A) Schematic structures of the two mismatch models hAmutsB-TCR and mutsAhB-TCR. (B) Schematic diagram of the plasmid sequences of the two mismatch constructs hAmutsB-TCR and mutsAhB-TCR. (C) Surface expression detection of the two mismatch constructs hAmutsB-TCR and mutsAhB-TCR in TCRαβ-KO Jurkat cells.

These results demonstrate that introducing engineered modifications while ablating the canonical disulfide bond enables TCRs to maintain high-efficiency surface expression and potent antitumor efficacy, while simultaneously minimizing mismatching. This design reduces mispairing-associated risks, thereby enhancing clinical applicability for future therapeutic implementation.

## Discussion

The efficacy of T cell therapies with engineered T cell receptors for solid tumors is limited by the problem of TCR mismatch. Although there are various optimization strategies for the TCR constant regions, either replacing the TCR constant regions with murine constant regions, adding additional disulfide bonds, or hydrophobic mutation of the TCR transmembrane structural domains have failed to completely solve the mismatch problem, and the efficacy of TCR-T needs to be further improved.

To address this, we pioneered an integrated approach by constructing sTCR incorporating three constant region optimizations (4857 + mm + TMa). While this achieved enhanced surface expression and cytotoxicity, the high surface expression of hA-sB-TCR and sA-hB-TCR heterodimers in TCRαβ-KO Jurkat cells revealed substantial mismatch risks.

We subsequently developed a novel solution targeting the canonical αCys95–βCys131 disulfide bond—the sole known mediator of endogenous TCR pairing. We deleted the endogenous intrinsic disulfide bond of sTCR by amino acid mutation, and the mutant mutEC-sTCR was still able to exhibit a high level of surface expression. The activation and cytotoxicity of mutEC-sTCR-engineered T cells were comparable to those of sTCR, which proved that disruption of canonical disulfide bond does not influence its assembly and functions. Although this optimized TCR has excellent tumor cytotoxicity *in vitro*, its *in vivo* safety profile and antitumor efficacy require further validation in animal models.

The mismatch models of mutEC-sTCR (hAmutsB-TCR, and mutsAhB-TCR) implied reduced exogenous-endogenous TCR mismatch. However, while β-chain mispairing was almost completely eliminated, optimized α-chain-still mediated a certain level of mismatching. What makes such a difference is still unknown. And the partial mismatch reduction suggests additional, undiscovered mechanisms may contribute to TCR interactions beyond the reported canonical disulfide bonds.

Our dual-strategy approach—retaining structural optimizations while ablating the canonical disulfide bond—significantly mitigates mispairing risks without compromising functionality. This represents a paradigm shift in TCR engineering. Based on this study, future work can be performed on (a)Engineering α-chain-specific solutions to eliminate residual mismatching, (b)Elucidating alternative chain-pairing mechanisms through structural studies, (c)Validating long-term safety and efficacy in NSG mouse tumor models.

Collectively, this study provides a critical foundation for clinical-grade TCR-T manufacturing with enhanced safety, advancing toward effective solid tumor immunotherapy.

